# Lipid phase separation in vesicles enhances TRAIL-mediated cytotoxicity

**DOI:** 10.1101/2021.10.29.466523

**Authors:** Timothy Vu, Justin A. Peruzzi, Lucas E. Sant’Anna, Neha P. Kamat

## Abstract

Ligand spatial presentation and density play important roles in many signaling pathways mediated by cell receptors and are critical parameters when designing protein-conjugated therapeutic nanoparticles. Currently, Janus particles are most often used to spatially control ligand conjugation, but the technological challenge of manufacturing Janus particles limits adoption for translational applications. Here, we demonstrate that lipid phase separation can be used to spatially control protein presentation onto lipid vesicles. We used this system to study the density dependence of TNF-related apoptosis inducing ligand (TRAIL), a model therapeutic protein that exhibits greater cytotoxicity to cancer cells when conjugated onto a vesicle surface than when administered as a soluble protein. Using assays for apoptosis and caspase activity, we show that phase separated TRAIL vesicles induced higher cytotoxicity to Jurkat cancer cells than uniformly-conjugated TRAIL vesicles, and enhanced cytotoxicity was dependent on the TRAIL domain density. We then assessed this relationship in other cancer cell lines and demonstrated that phase separated TRAIL vesicles only enhanced cytotoxicity through one TRAIL receptor, DR5, while another TRAIL receptor, DR4, was unaffected by the TRAIL density. These results indicate unique signaling requirements for each TRAIL receptor and how TRAIL therapy could be tailored depending on the relative levels of expression for cancer receptors of interest. Overall, this work demonstrates a readily adoptable method to control protein conjugation and density on bilayer vesicles that can be easily adopted to other therapeutic nanoparticle systems to improve receptor signaling of nanoparticles targeted to cancer and diseased cells.

## Introduction

The spatial presentation of ligands has a critical impact on cellular response. The density and inter-ligand distance of extracellular ligands interacting with their receptors can control binding and adhesion, ultimately affecting the initial steps of cell signaling pathways and controlling a range of processes from immune responses to stem cell differentiation.^1–3^ For ligand-conjugated therapeutic nanoparticles, the ligand density and spacing can affect cellular binding and uptake, *in vivo* pharmacokinetics, and receptor activation.^4,5^ By considering the role of ligand spatial presentation, therapeutic nanoparticles may be better designed to elicit desired cellular responses. Towards this goal, several Janus nanoparticles, particles with two unique surfaces that have distinct properties, have been developed to control the location and density of surface-conjugated ligands. These Janus nanoparticles have been shown to allow precise patterning of ligands in opposite faces, which have been used to study endocytosis of partial PEGylation,^6^ ligand clustering for T cell activation,^7,8^ and developing nanoparticles that mimic antibodies.^9^ Yet the assembly of these particles is often complex, requiring highly controlled chemistry and highly involved manufacturing schemes. The difficulty in creating Janus nanoparticles has hindered their adoption as therapeutic nanoparticles, despite their advantage of controlled ligand presentation.

In contrast to designed, colloidal nanoparticles, cellular membranes naturally control the spatial presentation of ligands using phase segregated lipid rafts. Lipid rafts are compartmentalized regions on the cell membrane characterized by increased cholesterol and saturated lipid content. These structures segregate distinct populations of biomolecules, affecting local membrane properties, lipid-protein interactions, and protein signaling.^10^ Phase separation can be readily replicated in synthetic, self-assembled lipid systems.^11^ Phospholipid bilayers, both in planar and nanoparticle form, can undergo phase separation when the appropriate mixtures of membrane components are used.^12,13^ Beyond a limited number of examples,^14,15^ this rich physical phenomenon has rarely been exploited to better design therapeutic nanoparticles.

A therapeutic target that could benefit from controlled presentation of ligands on a nanoparticle surface is the death receptor apoptosis pathway for anti-cancer therapy. In this pathway, TNF-related apoptosis inducing ligand (TRAIL) binds to death receptors DR4 and DR5, which are upregulated in cancer cells, and induces apoptosis.^16^ TRAIL naturally exists as a homotrimer where it exhibits optimal bioactivity. As functional TRAIL exhibits enhanced activity during oligomerization, recombinant TRAIL often requires further clustering or cross-linking for therapeutic use.^17^ For example, TRAIL signaling and anti-cancer activity have been shown to improve when conjugated onto a lipid vesicle (liposome) compared to soluble TRAIL.^18–20^ This improvement is observed because the vesicle membrane serves as a scaffold to promote TRAIL oligomerization on the cell surface, which is necessary for downstream signaling. Despite TRAIL improvements by conjugating to lipid vesicles, TRAIL still performs poorly *in vivo*. This is in part because we do not yet understand how the nature of TRAIL density impacts cell-signaling. Beyond liposome reconstitution, methods that allow for greater control of TRAIL presentation on nanoparticles should allow us to begin answering questions related to TRAIL-induced cell signaling and improve the anti-cancer activity of TRAIL-conjugated therapeutic nanoparticles.

We set out to test whether we could control TRAIL density through changes in lipid composition in vesicles, and whether the density of TRAIL could improve signaling and cancer cell apoptosis. By conjugating TRAIL to unsaturated lipids and varying the concentration of domain-forming saturated lipids, we reasoned that we could control TRAIL density in vesicles by decreasing the surface area available to TRAIL-conjugated lipids while keeping the concentration of TRAIL constant overall. Furthermore, we hypothesized that localization of TRAIL would improve receptor oligomerization on target cells and TRAIL-mediated signaling would increase, ultimately leading to higher levels of apoptosis. Using recombinant TRAIL, we show the extent to which lipid phase separation can be harnessed to change the spatial density of TRAIL on vesicle surfaces and reduce the concentration of TRAIL required to elicit cell death. By leveraging the intrinsic ability of lipid vesicles to phase separate, we demonstrate that lipid composition can be engineered to modulate the spatial density of proteins on a vesicle surface and promote TRAIL clustering and cytotoxic efficacy (**Scheme 1**).

**Scheme 1.**
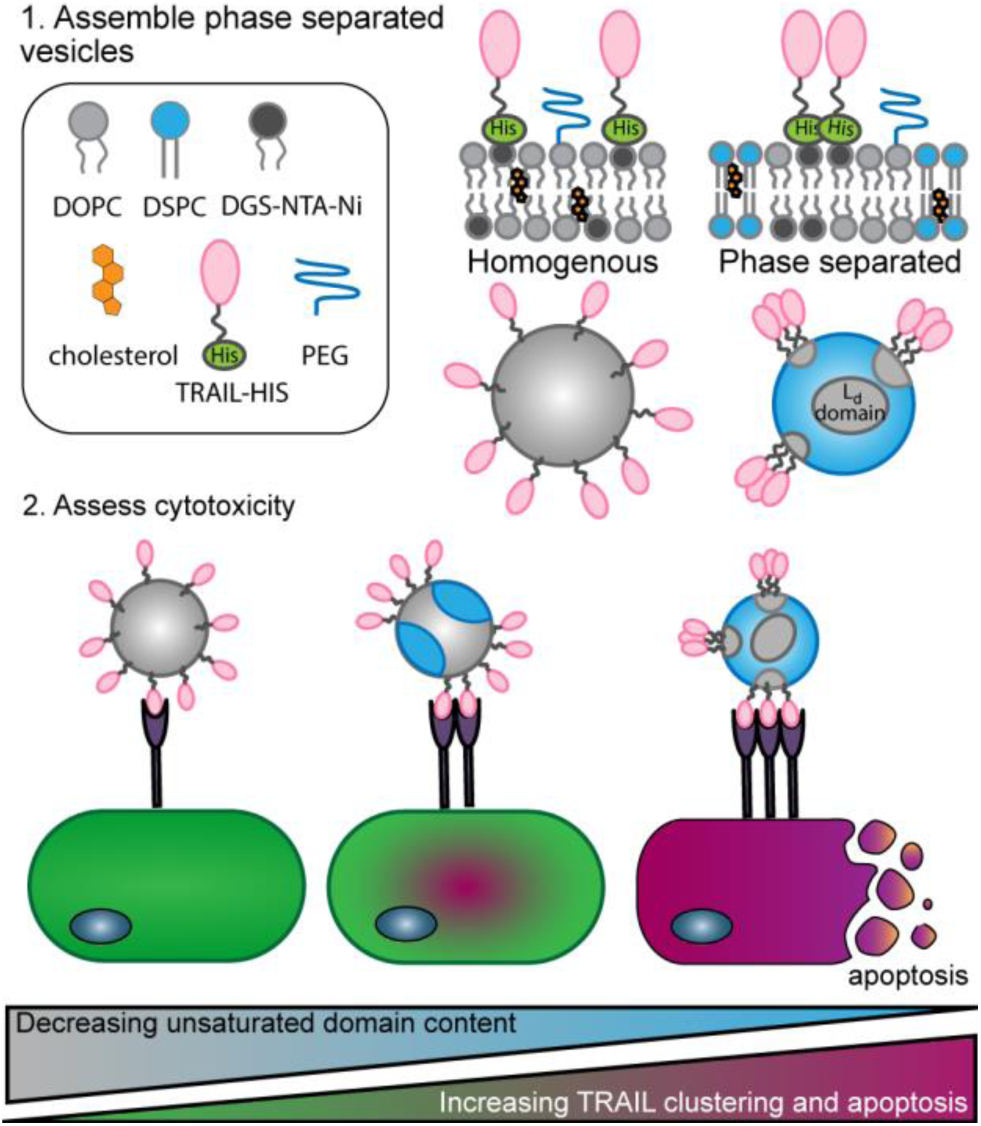
Design of TRAIL-conjugated nanoparticles using phase separated lipid vesicles. (Top) By varying the ratio of unsaturated lipid DOPC, saturated lipid DSPC, and cholesterol, lipid vesicles can be assembled containing homogenous or phase segregated membranes. A nickel-conjugated lipid (DGS-NTA-Ni) that integrates into the liquid disordered phase of the membrane can bind histidine-tagged TRAIL proteins and generate nanoparticles with varying spatial densities of TRAIL. Nanoparticles with low unsaturated lipid content and increasing density of TRAIL is hypothesized to enhance apoptosis in target cells.

## Results/Discussion

### Assembling phase separated lipid vesicles with surface-conjugated TRAIL

To control the surface density of TRAIL proteins on liposomes, we assembled nanovesicles with varying levels of membrane phase separation. DOPC (1,2-dioleoyl-sn-glycero-3-phosphocholine), an unsaturated lipid, forms a liquid-disordered (L_d_) phase, whereas DSPC (1,2-distearoyl-sn-glycero-3-phosphocholine)/cholesterol mixtures can form a liquid-ordered (L_o_) phase.^21^ When mixed at the appropriate ratios, these three lipids can phase separate into L_d_ and L_o_ phases within a single vesicle membrane. Lipid vesicles were assembled using ternary mixtures of DOPC, DSPC, and cholesterol at varying ratios. To localize TRAIL to select regions of the membrane, we included an unsaturated lipid with a Ni-NTA (nickel-nitrilotriacetic acid) headgroup that localized into L_d_ regions in the liposome membrane. This Ni-NTA lipid allowed conjugation of monomeric His-tagged TRAIL to our vesicles after vesicle formation. Polyethylene glycol (PEG, [18:0 PEG(2000)]), conjugated to a saturated lipid, was incorporated at 1 mol % for vesicle stability. We chose a PEG chain linked to a saturated lipid versus an unsaturated one because it should partition away from the Ni-NTA-conjugated lipids and reduce possible steric hindrance of TRAIL. Vesicles were extruded to 100 nm, incubated with His-tagged TRAIL, and dialyzed to remove unconjugated protein. By increasing the ratio of DSPC to DOPC in these vesicles, we expected to increase the size of L_o_ domains and reduce the size of L_d_ domains, respectively.

We used both microscopy and Förster resonance energy transfer (FRET) analysis to confirm lipid phase separation. Vesicles were labeled with a saturated lipid dye (16:0 NBD) and an unsaturated lipid dye (18:1 Rho, [Lissamine-Rhodamine PE]) that should separate into L_o_ and L_d_ domains, respectively. Microscopy of giant unilamellar vesicles (GUVs) formed through electroformation^11^ showed segregation of 16:0 NBD and 18:1 Rho into distinct regions of the membrane in ternary compositions of DOPC, DSPC and cholesterol, confirming the formation of phase-separated lipid domains (Figure 1A). In compositions with only DOPC or DSPC, the two dyes mixed well and did not segregate. Next, we measured the FRET efficiency between 16:0 NBD (donor) and 18:1 Rho (acceptor) to assess the spatial separation between L_o_ and L_d_ phases in nano-scale vesicles composed of differing ratios of DOPC and DSPC as previously described (Figure 1B).^22,23^ These studies demonstrated measurable differences in FRET ratio (here reported as normalized F_donor_/F_acceptor_ [see Equation 1 and 2]) as a function of DSPC content, suggesting differences in phase separation as a function of vesicle composition (Figure 1C). As expected, vesicles composed of DOPC alone showed the lowest FRET ratio, indicating the two fluorescent probes were uniformly distributed across the membrane. Increases in the mole fraction of DSPC increased the FRET ratio, indicating the saturated and unsaturated lipids were located farther apart within a given membrane and suggesting the presence of nanodomains. Pure DSPC vesicles exhibited the highest FRET ratio, which we attribute to the presence of extremely small L_d_ domains that result from segregation of the NTA-lipids and unsaturated lipid dye. The conjugation of TRAIL to NTA lipids did not change FRET values of all vesicle compositions relative to their FRET values prior to TRAIL conjugation, indicating the conjugation of TRAIL did not cause domain dissociation (Figure 1D). To further confirm that the FRET data reflected the presence of membrane domains, we used a temperature ramp to destabilize domains.^11,22^ Our temperature studies show that the FRET ratio is temperature-dependent and decreased at higher temperatures; this result is expected, as domains are known to dissolve and lipids become well mixed when approaching the melting temperature (T_m_) of the saturated lipid (55°C for DSPC, Supporting Figure S1). In summary, our characterization of giant and nano-sized vesicles confirms that our choice of lipid mixtures yields vesicles with varied degrees of lipid segregation, a spatial organization that is conserved upon TRAIL conjugation.

**Figure 1.**
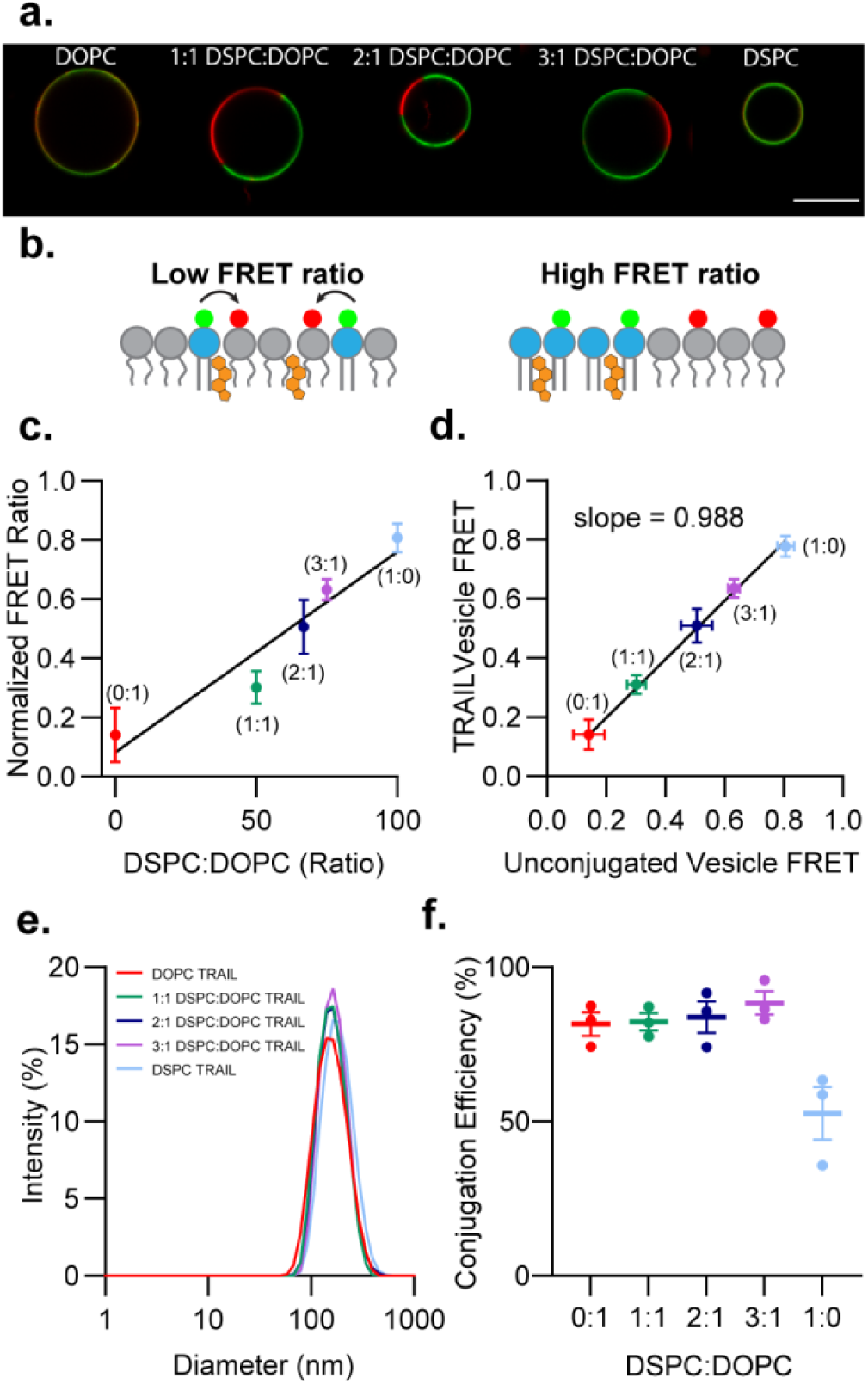
Characterization of TRAIL conjugation to lipid domain vesicles. (a) Microscopy images of GUVs to visualize phase separation of saturated lipids (green) with unsaturated lipids (red). Scale bar = 10 μm. (b) Schematic of a FRET assay to determine lipid domain presence in vesicles. (c) FRET analysis of vesicle lipid domains before TRAIL conjugation, where increasing FRET Ratio indicates the presence of domains. FRET ratio is reported as F_donor_/F_acceptor_. Error bars represent SEM from n = 3 different vesicle preparations. (d) FRET analysis of vesicle lipid domains before and after TRAIL conjugation shows no change after TRAIL conjugation. Error bars represent SEM from n = 3 different vesicle preparations. (e) Size distribution of TRAIL-conjugation vesicles measured by dynamic light scattering (DLS). (e) Conjugation efficiency of TRAIL to vesicles, as determined by western blot. Error bars represent SEM from n = 3 different vesicle preparations and Western Blots.

Next, we measured several physical and chemical properties of TRAIL-conjugated vesicles to ensure vesicles were similar in size and total concentration of TRAIL per vesicle. All TRAIL-conjugated vesicle compositions exhibited similar size and zeta potential (Figure 1E and Supporting Table S2). We next confirmed the extent of TRAIL conjugation onto the vesicles using western blot analysis. DOPC, 1:1 DSPC:DOPC, 2:1 DSPC:DOPC, and 3:1 DSPC:DOPC compositions all showed similar total levels of TRAIL conjugation, while, surprisingly, DSPC vesicles demonstrated less TRAIL conjugation (Figure 1F and Supporting Figure S3A). We hypothesize that TRAIL conjugation to DSPC vesicles is poor because the conjugating lipid (DGS-NTA-Ni) is unsaturated and assembles into extremely small L_d_ domains. As a result, steric hindrance from conjugated TRAIL molecules could prevent other TRAIL molecules from associating with unoccupied/unbound DGS-NTA-Ni. Non-reducing western blot analysis revealed oligomeric TRAIL structures in vesicles with domains, with more oligomeric structures as the L_o_ domain size decreased (Figure S3B). DSPC vesicles exhibited extremely large oligomers, which supports our hypothesis that TRAIL-conjugated lipids are highly segregated in this region and likely to prevent other TRAIL molecules from binding to unbound DGS-NTA-Ni. Nonetheless, all other compositions showed similar levels of TRAIL conjugation. We therefore established a series of membrane compositions that displayed differential phase separation but similar overall total TRAIL concentrations that would allow us to isolate the role of TRAIL spatial density on cell cytotoxicity.

### TRAIL conjugation to phase segregated lipids enhances TRAIL-mediated apoptosis

Next, we studied the capacity of TRAIL-conjugated vesicles to initiate TRAIL-mediated apoptosis. To measure cytotoxicity of TRAIL-conjugated vesicles, we chose to first study Jurkat cells. This cell type is more sensitive to TRAIL-conjugated vesicles than soluble TRAIL alone, which suggests Jurkat cells might be responsive to further spatial localization of TRAIL molecules within membrane domains.^20^ Jurkat cells were treated with increasing concentrations of either soluble TRAIL or TRAIL-conjugated vesicles for 24 hours (Figure 2A). Despite DOPC TRAIL and soluble TRAIL possessing similar relative IC_50_ values (29 ng/mL and 25 ng/mL, respectively), DOPC TRAIL vesicles were more efficacious, reducing viability to approximately 25% while the effect of soluble TRAIL plateaued at 50% even at higher concentrations (Supporting Figure S4). Furthermore, we observed that segregation in vesicle domains increased the cytotoxicity of TRAIL and that cytotoxicity correlated with smaller domain size (3:1 DSPC:DOPC > 2:1 DSPC:DOPC > 1:1 DSPC:DOPC > DOPC). Decreasing the domain size both increased the efficacy, as domain vesicles killed virtually all Jurkat cells at 200 ng/mL, and potency (relative IC_50_ of 17 ng/mL for 1:1 DSPC:DOPC, 7 ng/mL for 2:1 DSPC:DOPC, and 6 ng/mL for 3:1 DSPC:DOPC). Surprisingly, DSPC vesicles did not exhibit any significant TRAIL-mediated cytotoxicity. While DSPC TRAIL vesicles had significantly less TRAIL conjugated on their surface, the concentration of TRAIL on DSPC vesicles (approximately 100 ng/mL at highest dose treated) should still be sufficient to induce apoptosis and indicates some other feature of TRAIL presentation in these particles is likely affecting their efficacy. To rule out the possibility that the cytotoxic effects we observed were due to the lipid vesicles themselves, we treated Jurkat cells with unconjugated vesicles at the highest lipid concentration used (Figure 2B). As expected, Jurkat viability was not negatively affected by unconjugated vesicles, though we did notice a slight increase in Jurkat growth with addition of DSPC vesicles. Binding studies of the vesicles to Jurkat cells demonstrated that DSPC increased binding of vesicles to cells nonspecifically, and that binding did not correlate with TRAIL-mediated cytotoxicity (Supporting Figure S5). We hypothesize that the reduced cytotoxicity of DSPC-TRAIL vesicles may be due to the increased rigidity of DSPC vesicles or an altered presentation of TRAIL molecules on the surface of DSPC vesicles. This is supported by previous studies that demonstrated membrane fluidity affected immunoliposome binding towards target cells.^24,25^ Our other vesicle compositions that showed robust TRAIL activity contained more unsaturated lipids and correspondingly more fluid membranes. Altogether, these results indicate that clustering TRAIL within L_d_ domains of lipid vesicles increases cell signaling and apoptosis.

**Figure 2.**
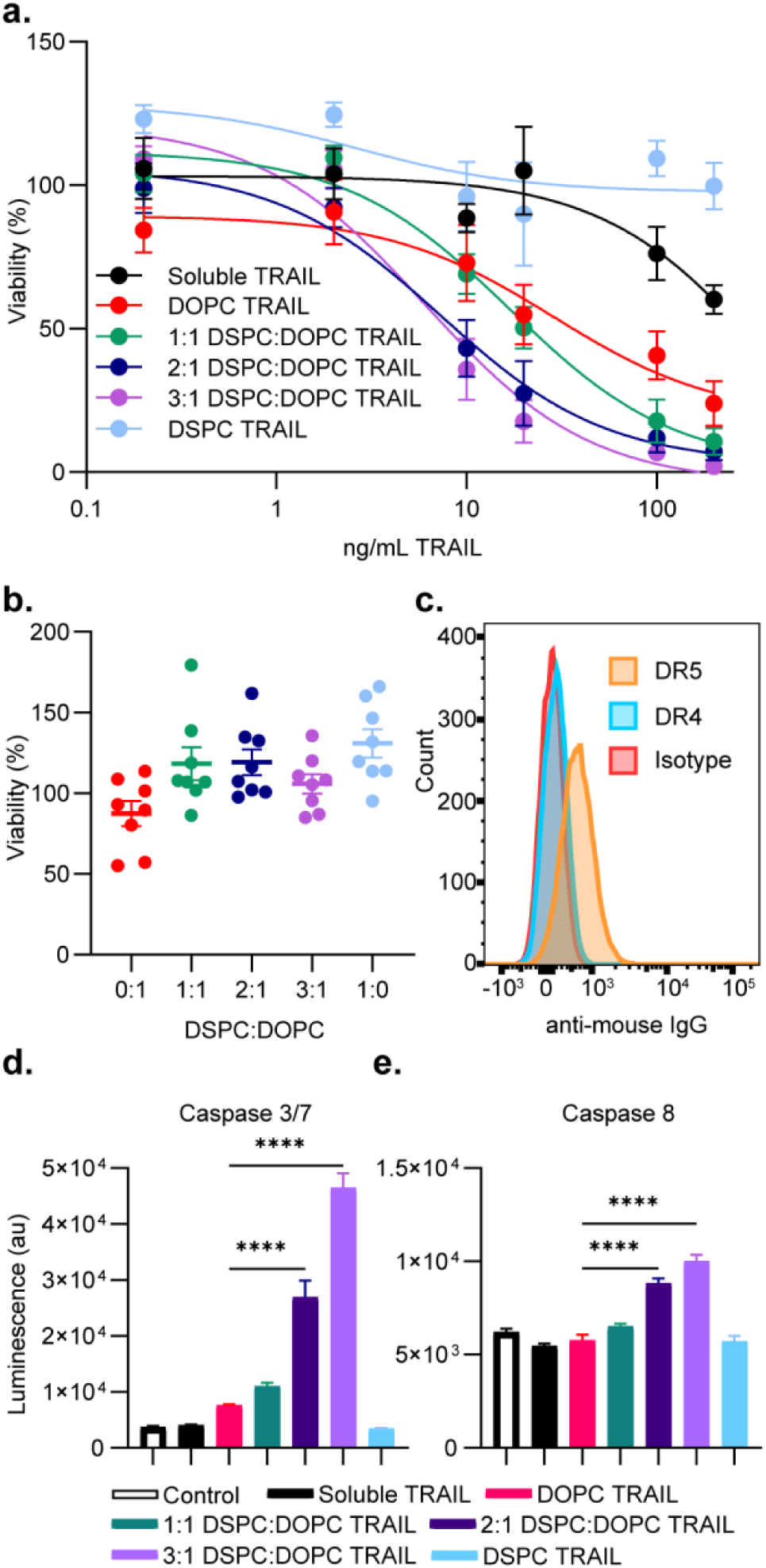
Vesicle lipid domains enhance TRAIL activation in Jurkat cells. (a) Viability of Jurkat cells treated with different concentrations of soluble and vesicle TRAIL after 24 hours. Error bars represent SEM from n = 6 using 2 different vesicle preparations. Concentration reported is the initial amount of TRAIL added to the vesicles during conjugation. (b) Viability of Jurkat cells to 1 mM unconjugated lipid vesicles, which corresponds to highest TRAIL vesicle concentration tested. Error bars represent SEM from n = 8 from 3 different vesicle preparations. (c) Flow cytometry analysis of Jurkat expression of TRAIL receptors DR4 and DR5. (d,e) Activation of caspase 3/7 (d) and caspase 8 (e) in Jurkat cells exposed to soluble and vesicle TRAIL (20 nM) after 3 hours. Error bars represent SEM from n = 3. Significance test used was ANOVA. **** p < 0.0001, *** p < 0.001, ** p < 0.01, * p < 0.05.

To further confirm TRAIL-mediated cell death, we next investigated the activation of intracellular apoptotic signaling molecules by monitoring caspase-3/7 and caspase-8 activity. When TRAIL binds to its receptors, death receptors 4 and 5 (DR4, DR5), it recruits FAS-associated protein with death domain (FADD) and pro-caspase-8 to the cell membrane. This leads to caspase-8 activation and subsequent activation of caspase-3 and caspase-7 to induce apoptosis.^26^ We therefore measured the activity of caspase-3/7 and caspase-8 to confirm cell death was the result of TRAIL-induced intracellular signaling. We measured the caspase-3/7 and caspase 8 activity in Jurkat cells after 3 hours of incubation with vesicles containing 20 ng/mL of TRAIL. (Figure 2C and 2D, respectively). At this time point and concentration we saw minimal caspase activity from soluble TRAIL, DOPC TRAIL, 1:1 DSPC:DOPC TRAIL or DSPC TRAIL, but vesicles with smaller domains in the 2:1 DSPC:DOPC TRAIL and 3:1 DSPC:DOPC TRAIL compositions showed significantly increased caspase activity. Therefore, segregation of TRAIL in the L_d_ domain of vesicles increased caspase 3/7 and caspase 8 activity after 3 hours over DOPC vesicles presenting uniformly distributed TRAIL. Curiously, DOPC TRAIL did not show any differences in caspase activity relative to soluble TRAIL at this time point despite increased cytotoxic activity over soluble TRAIL after 24 hours, which could indicate that clustering also affects the rate of caspase activity. Work studying Fas/FasL, which induces apoptosis similarly to TRAIL/DR5, demonstrated the ligand spacing of FasL affected the apoptosis rate, which could explain increased caspase activity in the phase segregated 2:1 and 3:1 DSPC:DOPC TRAIL compositions but not DOPC TRAIL.^27^ These results confirmed the mode of TRAIL-vesicle mediated cytotoxicity was through caspase-mediated apoptosis, and that concentrating TRAIL into vesicle membrane domains increased caspase activity.

Finally, we wanted to assess the reproducibility of our system with other cell types. We assessed the cytotoxicity of phase segregated, TRAIL vesicles on 5 additional cancer cell lines: U2-OS bone osteosarcoma, U937 monocytic leukemia, MDA-MB-231 triple negative breast adenocarcinoma, K562 bone marrow leukemia, and HCT-116 colon colorectal carcinoma (Figure 3A-E). Interestingly, we found that TRAIL-segregation in membrane domains did not universally improve vesicle cytotoxicity for all cancer lines tested. Some cell types were sensitive to TRAIL segregation in domains similar to Jurkat cells (U2-OS, U937), while other cell types were showed no differences in viability between soluble TRAIL or domain-segregated TRAIL (K562, KDA-MB-231, and HCT-116). Again, treatment of cells with unconjugated vesicles at the highest lipid concentration showed no effects on viability, suggesting all observed cell death was dependent on TRAIL interactions with cells (Supporting Figure S6). DSPC TRAIL vesicles again showed limited cytotoxicity to all cell lines except HCT-116 (Supporting Figure S7). We further tested caspase activity in U937 cells, one of the cells affected by domains, and saw similar trends to what was observed in Jurkat cells (Supporting Figure S8). These results indicate that the capacity of spatial segregation to improve TRAIL mediated cell death is dependent on cell type.

**Figure 3.**
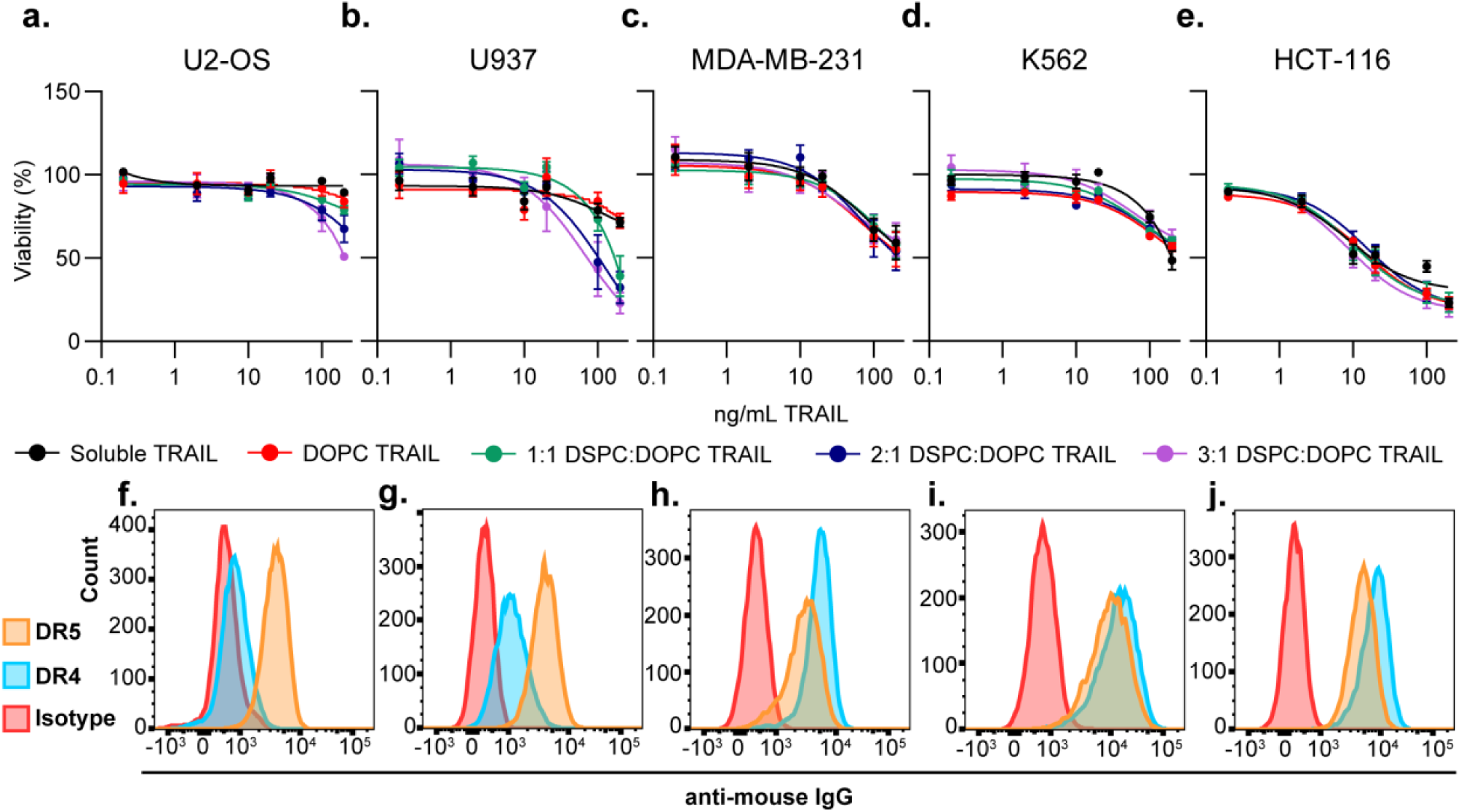
Differential effects of TRAIL vesicle lipid domains on various cell lines. Viability treated with soluble and vesicle TRAIL after 24 hours and expression of TRAIL receptors DR4/DR5 of U2-OS (a,f), U937 (b,g), MDA-MB-231 (c,h), K562 (d,i) and HCT-116 (e,j). Error bars represent SEM from n = 6 from two different vesicle preparations. Concentration reported is the initial amount of TRAIL added to the vesicles during conjugation.

### DR5/DR4 expression affects the impact of phase segregated TRAIL vesicles

We wondered why certain cell types demonstrated higher sensitivity to TRAIL segregation versus other cells. We hypothesized that vesicle interactions with TRAIL receptors DR4 and DR5 may play a role in TRAIL potency. Using flow cytometry, we measured the expression levels of DR4 and DR5 for each cell type (Figure 3F-J). We then examined which cell types were sensitive to TRAIL segregation on our vesicle formulations. We compared the difference in cell viability between cells incubated with TRAIL conjugated to 3:1 DSPC:DOPC or DOPC vesicles, where TRAIL was segregated or uniformly distributed, respectively, and identified two distinct clusters (Figure 4). Cells that predominantly expressed DR5 over DR4 were sensitive to TRAIL segregation in vesicle domains, while cells that expressed more or equivalent amounts of DR4 over DR5 showed no viability differences as a function of vesicle composition.

**Figure 4.**
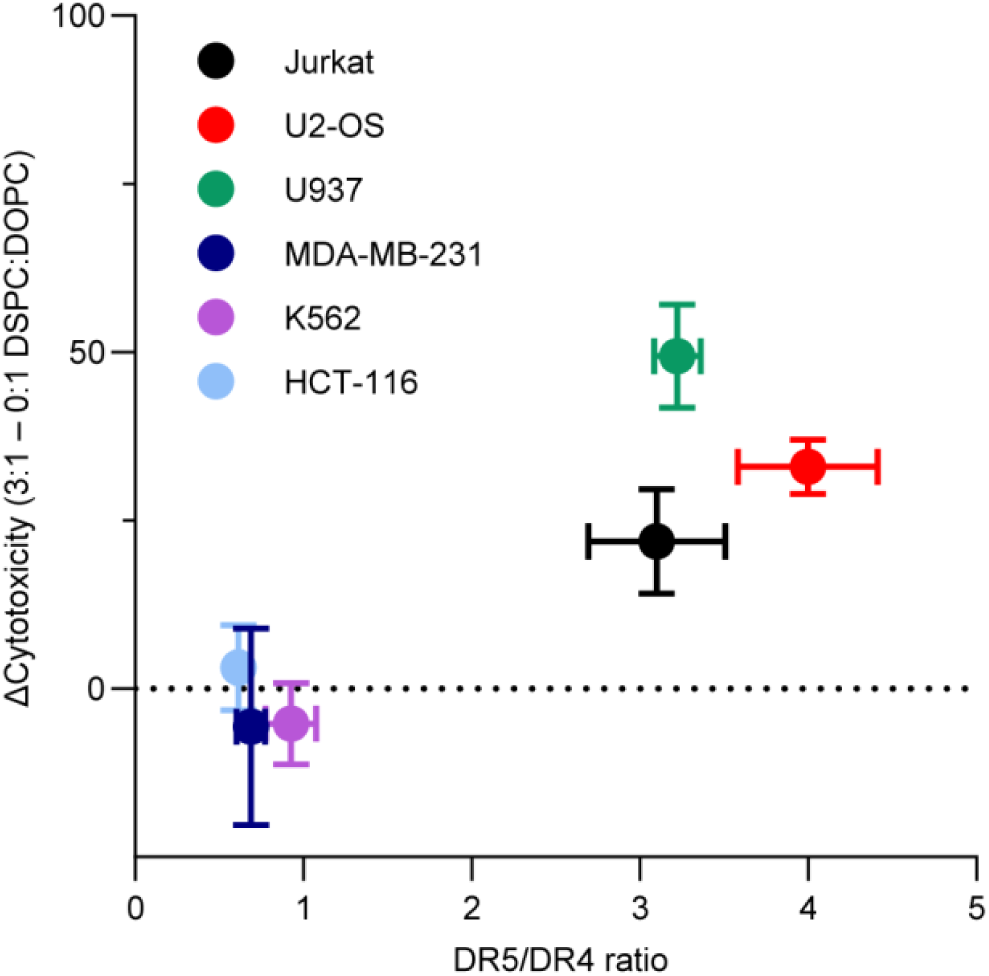
TRAIL domain enhancement is dependent on DR4 and DR5 expression levels. X-axis is the DR5/DR4 ratio, and Y-axis is the cytotoxicity difference between 3:1 DSPC:DOPC TRAIL vesicles to pure DOPC TRAIL vesicles. Cytotoxicity differences between domains and DOPC vesicles are only seen when DR5 is expressed higher than DR4. Error bar represents the SEM from n = 6 (y-axis) and n = 3 (x-axis).

Previous studies have shown that DR5 and DR4, which must oligomerize into supramolecular structures to initiate apoptosis, can respond differentially to TRAIL structure. Specifically, DR5 benefits from oligomerization or cross-linking of TRAIL in order to activate apoptotic signaling.^28–30^ In contrast, the aggregation state of TRAIL has much less effect on DR4 activation.^28,31,32^ TRAIL density may have less of an effect on DR4 activation in part because DR4 resides in lipid rafts in certain cell types that are susceptible to TRAIL.^33^ The lipid raft, in essence, could pre-organize the receptors and benefit less from the organization of TRAIL ligands. Because DR4 can competitively bind TRAIL in the presence of DR5,^31,34,35^ the relative differences in receptor expression should dictate the extent to which TRAIL density influences apoptotic signaling. We expect, therefore, that when DR5 is more prevalent on cell membranes than DR4, the density of TRAIL should become more important, and vesicles that localize TRAIL in phase segregated domains should affect the extent of apoptotic signaling. In contrast, when DR4 is more prevalent and TRAIL signaling occurs primarily through this receptor, the spatial density of TRAIL should be less important. This was observed in our results with TRAIL domains against different cell types with different DR4 and DR5 expressions. Besides competitive binding, DR4 and DR5 can also form heterocomplexes, which is not well understood but has been shown to predominantly signal through DR4 in pancreatic tumor cells.^36,37^ Ultimately, relative levels of DR4 and DR5 expression, and which receptor the cell predominately signals through, could affect the therapeutic strategy for TRAIL agonists.

Lipid domains provide a straightforward route to control the spatial density of a lipid-linked protein to signal cluster-dependent receptors like DR5. Our observations of TRAIL mediated cytotoxicity in cell types that express higher levels of DR5 relative to DR4 show that concentrating TRAIL in smaller lipid domains enhances cytotoxicity relative to vesicles with homogeneously distributed TRAIL. This observation is consistent with recent studies using DNA origami nanostructures in which precise ligand spatial orientation of DR5 improved apoptosis, and that an inter-ligand distance of approximately 5 – 10 nm was found to be optimal for DR5-mediated cell death.^38^ A similar study found precise ligand spatial orientation was critical for Fas (CD95) signaling, another TNF superfamily receptor with a similar signaling pathway as DR5.^27^ We hypothesize that controlling TRAIL spatial orientation on lipid vesicles by using lipid domains most likely increases DR5-mediated apoptosis by approaching this critical inter-ligand distance. Taken together, lipid phase separation is a facile method to control the spatial orientation of lipid linked proteins that can be readily incorporated into lipid nanoparticle design.

## Conclusion

The physical features of therapeutic nanoparticles have a critical impact on their efficacy. Particle size, shape, charge, and stiffness, for example, are all well-known parameters that dictate cellular uptake, endosomal escape, cellular signaling, and ultimately therapeutic efficacy.^39^ Among these properties, the spatial presentation of surface-conjugated molecules has been less explored. In our current study, we have demonstrated that phase separation of lipids into L_o_ and L_d_ domains on vesicles can be used to spatially control protein conjugation. By controlling the spatial conjugation of TRAIL into L_d_ domains of different sizes to increase the local spatial density, we demonstrate increased efficacy of TRAIL-mediated apoptosis in cancer cells. Increases in TRAIL-mediated apoptosis were dependent on the size of the L_d_ domain, with the smallest L_d_ domains on DOPC:DSPC ternary vesicles exhibiting the greatest TRAIL-mediated apoptosis and caspase activity. The fluidity and structure of the vesicle also appear to play a role, as predominantly DSPC vesicles, which exhibited the smallest L_d_ domains, were less effective at facilitating cell death relative to the other nanoparticle compositions. Additionally, increases in cell death based on spatial conjugation of TRAIL was dependent on DR5 signaling; spatial conjugation of TRAIL into L_d_ domains of vesicles increased cytotoxicity to cells that expressed greater levels of DR5 compared to DR4, while cells that expressed similar levels of DR5 and DR4 showed no differences in cell death when treated with vesicles exhibiting TRAIL domains. Spatial presentation of ligands on therapeutic nanoparticles therefore must be optimized towards the biology of the target receptor.

Other types of nanoparticle approaches have been used to enhance anti-tumor activities of TRAIL. For example, TRAIL has been conjugated onto non-spherical virus particles^26^ and carbon nanotubes^40^ to enhance apoptosis. Polymeric nanoparticles have acted as “mechanical amplifiers” to sensitize TRAIL.^41^ TRAIL nanoparticles can be attached to many different types of blood-circulating cells to increase the efficacy of *in vivo* delivery.^42–44^ A gap in TRAIL-mediated material therapies, however, has been to understand how optimize the density of TRAIL to maximize apoptosis. As our approach can be readily incorporated into the aforementioned nanoparticles or cellular designs, we foresee that including investigations into TRAIL surface density for these and other nanoparticle-based therapies would help further improve TRAIL efficacy for cancer therapy and push TRAIL to success in the clinic. Phase separated vesicles are simple to create, and highly biocompatible, which is an important technological advantage over Janus particles and will allow for a greater degree of freedom when developing therapeutic nanoparticles.

Finally, vesicle phase separation is a useful technique to enhance protein-conjugated vesicles beyond TRAIL liposomes. From a biomaterials perspective, controlled spatial conjugation of proteins can apply to other types of vesicles with demonstrated phase separation, such as lipid-polymer hybrid vesicles^23,45^ and polymersomes.^46,47^ as well as in supported bilayer systems for fundamental investigations into the effects of spatial arrangements on biological mechanisms.^48^ As such, our work describes a technique to control ligand density that can be applied to a wide variety of nanoparticle types and biological studies. Biological systems that are also known to be dependent on receptor clustering include immune signaling receptors^49–51^ and receptor internalization.^52,53^ Altogether, phase separated vesicles provide researchers a new tool to spatially control protein spacing for designing cell-mimetic systems and therapeutic nanoparticles.

### Experimental Section

#### Materials

1,2-dioleoyl-sn-glycero-3-phosphocholine (DOPC), 1,2-distearoyl-sn-glycero-3-phosphocholine (DSPC), cholesterol (Chol), 1,2-dioleoyl-sn-glycero-3-[(N-(5-amino-1-carboxypentyl)iminodiacetic acid)succinyl] (nickel salt) (DGS-NTA-Ni), 1,2-distearoyl-sn-glycero-3-phosphoethanolamine-N-[methoxy(polyethylene glycol)-2000] (18:0 PEG2000-PE), 1,2-dioleoyl-sn-glycero-3-phosphoethanolamine-N-(lissamine rhodamine B sulfonyl) (18:1 Rho) were purchased from Avanti Polar Lipids. 1,2-dipalmitoyl-sn-glycero-3-phosphoethanolamine-N-(7-nitro-2-1,3-benzoxadiazol-4-yl) (16:0 NBD) was purchased from Invitrogen. Recombinant human TRAIL (His-tagged) was purchased from R&D Systems. CellTiter-Glo 2.0, Caspase-Glo 3/7, and Caspase-Glo 8 assay kits were purchased from Promega. Phosphate-buffered Saline (PBS) tablets were obtained from Sigma-Aldrich. Cell media components DMEM, RPMI, McCoy’s 5A, fetal bovine serum (FBS) and penicillin streptomycin were purchased from Thermo Fisher (Gibco). Antibodies used for flow cytometry were anti-DR4 (clone HS101, Novus Biologicals), anti-DR5 (clone HS201, Novus Biologicals), isotype control (biotinylated clone MOPC-21, Biolegend), anti-mouse IgG (Cell Signaling Technology 7076) and APC-labeled anti-mouse IgG (Ab510115, Abcam).

#### Vesicle Formation

Small unilamellar vesicles (SUVs) were prepared using the thin film hydration method. Lipids dissolved in chloroform were dried down under a nitrogen stream to create lipid films and placed in a vacuum for 2 hours. Lipid films were rehydrated overnight with 1x PBS (290 mOsm, pH 7.3) at 60°C. Vesicles were briefly vortexed and extruded using an Avanti mini-extruder through a 100 nm polycarbonate filter placed on a hot plate at 70°C for nine passes. Lipid compositions for each sample are provided in the supplement. Giant unilamellar vesicles (GUVs) were formed through electroformation using a Vesicle Prep Pro (Nanion).

#### FRET assays

Vesicles containing 0.5 mol% of 18:1 Rho (acceptor) and 0.5 mol% 16:0 NBD (donor) were added to a cuvette and the fluorescence was measured at 460 nm excitation and 535 nm and 583 nm emission using an Agilent Cary Eclipse Fluorimeter at 37°C. 0.1% Triton-X was added to each cuvette to lyse the vesicles to measure the unquenched fluorescence. For temperature ramp experiments, fluorescence was measured from 25°C to 58°C at 1°C steps.

FRET ratio and Normalized FRET ratio are defined as:

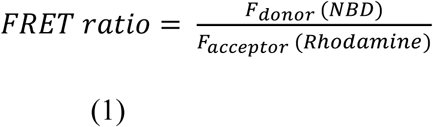

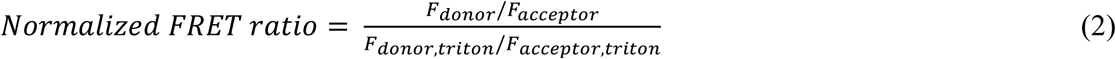

#### TRAIL conjugation to vesicles

To conjugate His-TRAIL onto vesicles, 1.0 mol% DGS-NTA lipid was incorporated into the vesicles to react with His-tag proteins, while 1.0 mol% 18:0 PEG2000-PE was added for vesicle stability. Vesicles were incubated with His-TRAIL (1 mM vesicles, 400 ug/mL TRAIL final concentration) for 1 hour at 37°C, then dialyzed overnight at 4°C with a 100 kDa dialysis kit (Float-A-Lyzer G2, Repligen). Size and zeta potential of vesicles after TRAIL conjugation were measured using a Malvern Zetasizer. For FRET studies after TRAIL conjugation, 0.5 mol% of 18:1 Rho and 0.5 mol% 16:0 NBD were included in vesicle samples prior to TRAIL conjugation and FRET efficiency was measured as previously described.

#### Western blot analysis

Efficiency of TRAIL conjugation was assessed by Western Blot. Briefly, samples were mixed with Laemmli buffer (after purification to remove unconjugated TRAIL for vesicles) and boiled at 95°C for 10 minutes (with and without SDS for reducing and non-reducing western blots, respectively). Samples were then loaded onto a 12% Mini-PROTEAN TGX Precast Protein Gel (Bio-Rad) and run at 100 V for at least 90 minutes. Wet transfer was performed onto an Immuno-Blot PVDF membrane (Bio-Rad) for 45 min at 100 V. Membranes were then blocked at room temperature for 1 hour in 5% milk in Tris-buffered saline with Tween 20 (TBST) pH 7.6 (20 mM Tris base, 150 mM NaCl, 0.1% Tween 20). Membranes were incubated in primary antibody solution (Mouse-anti-TRAIL (R&D Systems, MAB687), diluted 1:1000 in 5% milk in TBST) overnight at 4°C. Primary antibody was decanted and membrane was washed three times with TBST pH 7.6. Membranes were then incubated in secondary antibody solution (HRP-anti-Mouse (CST 7076), diluted 1:3000 in 5% milk in TBST pH 7.6) for 1 hour at room temperature. After incubation with secondary solution antibody, membranes were washed 3 times in TBST pH 7.6 and incubated with Clarity Western ECL Substrate (Bio-Rad) for 5 min and then imaged with an Azure C280. Images were then analyzed using the Fiji gel analysis tool.^54^

#### Cell culture

Jurkat, U937, HCT-116, and U2OS cells were obtained from ATCC without further authentication. MDA-MB-231 was obtained as a gift from the Mrksich lab from ATCC (Northwestern University) and K562 was obtained as a gift from the Leonard lab from ATCC (Northwestern University). Jurkat, U937, and K562 were cultured in RPMI 1640 supplemented with 10% FBS and 1% Pen/Strep. U2OS and HCT-116 were cultured in McCoy’s 5A supplemented with 10% FBS and 1% Pen/Strep. MDA-MB-231 were cultured in DMEM supplemented with 10% FBS and 1% Pen/Strep.

#### TRAIL cell viability assays

For adherent cells (MDA-MB-231, HCT-116, and U2OS), cells were detached from the plate using TrypLE Express and seeded in 96-black clear-bottom well plates at 5,000 cells/well the day before TRAIL addition to allow the cells to adhere. Suspension cells (Jurkat, U937, K562) were seeded at 25,000 cells/well the day of TRAIL addition. Vesicles in PBS and media (50% full media, 50% vesicles in PBS, 100 uL final volume) were incubated with cells at the indicated concentration (TRAIL is assumed to be 100% conjugated on vesicles when compared to soluble TRAIL) for 24 hours. After 24 hours, 100 uL of Celltiter Glo 2.0 solution was added to each well, allowed to mix for 5 minutes, then the luminescence was read with a plate reader (Molecular Devices). To analyze the percent cell viability, the luminescence of the treated well was divided by the luminescence of untreated cell wells and converted to a percentage. Viability studies were performed with at least two different vesicle samples in triplicate.

#### Vesicle binding studies to Jurkat

Jurkat cells were seeded at 100,000 cells/well in a 96 U-bottom plate in flow buffer (1% FBS in PBS). Vesicles containing 0.5% 18:1 Rho at the indicated concentrations were incubated with Jurkat cells for one hour at room temperature (RT), washed two times, and resuspended in 200 μL of flow buffer. Cells were analyzed using a BD Fortessa LSRII using 550nm excitation laser and 582/15 nm emission. Jurkat cells were gated based on forward vs side scatter (FSC-A vs. SSC-A) and single cells (FSC-A vs. FSC-H) and 10,000 single cell events were collected. Data was analyzed using FlowJo. Experiments were repeated in triplicate.

#### TRAIL caspase assays

Jurkat and U937 cells were seeded at 25,000 cells/well and 20 ng/mL soluble TRAIL or vesicle TRAIL was added to a 96-black clear-bottom well plate. Cells and TRAIL were incubated for 3 hours, then Caspase 3/7 and Caspase 8 detection kits (Promega) were added according to the manufacturer’s protocols. Luminescence was read at every 5 minutes for 1 hour, and the max signal was used. Luminescence was normalized to the untreated cells.

#### DR4 and DR5 expression

DR4 and DR5 expression of cells was measured using flow cytometry. For Jurkat, MDA-MB-231, U2-OS, and HCT-116, cells were incubated with anti-DR4, anti-DR5, or isotype control at 1 μg/mL in flow buffer for 30 minutes at RT, washed, then 2 μg/mL of APC anti-mouse IgG was added for 30 minutes at RT. Cells were washed 2x, resuspended in flow buffer, then analyzed using BD Fortessa LSRII using 640nm excitation and 670/14 nm emission. Cells were gated as previously described, and 10,000 single cell events were collected. Data was analyzed using FlowJo. For U937 and K562 cells, a modified protocol was used because of nonspecific binding of the isotype control. Cells were first incubated with isotype control in flow buffer for 30 minutes on ice, washed, then blocked with anti-mouse IgG in flow buffer for 30 minutes on ice. Blocked cells were untreated or incubated with anti-DR4 and anti-DR5 for 30 minutes at RT, washed, then incubated with 2 μg/mL of APC anti-mouse IgG for 30 minutes at RT. Cells were washed 2x and analyzed similarly to the other cells. Expression studies were done in triplicate, and the reported histogram expression data is a representation of the triplicate experiments.

#### Statistical Analysis

All nonlinear fits (three parameters), IC_50_ values, and significance tests were performed using Prism (Graphpad).

## Acknowledgements

This work was supported in part by the Searle Funds at The Chicago Community Trust and the National Science Foundation under Grant No. 1844219.. T.Q.V. was supported by the National Institutes of Health Training Grant (T32GM008449) through Northwestern University’s Biotechnology Training Program. J.A.P. gratefully acknowledges support from the Ryan Fellowship and the International Institute for Nanotechnology at Northwestern University. J. A. P. was supported by an NSF Graduate Research Fellowship. This work made use of the Northwestern RHLCCC Flow Cytometry Facility (NCI CA060553) and Northwestern NUANCE center (NSF ECCS-2025633 & NSF DMR-1720139). We thank the Kamat lab members for careful reading of the manuscript and the Mrksich group for helpful discussions.

## Author contributions

T.Q.V., J.A.P., and N.P.K. conceived the study and designed experiments. T.Q.V., J.A.P., and L.E.S. performed the experiments, T.Q.V., J.A.P., and N.P.K. analyzed the data. All the authors contributed to writing the manuscript.

