## Supporting Information for "Lipid phase separation in vesicles enhances TRAIL-mediated cytotoxicity"

**List of Supporting Information**

**Figure S1.** Temperature dependence of lipid domains measured by FRET ratio

**Table S2.** Summary values of size, polydispersity (PDI), and Zeta potential of TRAIL-conjugated vesicles

**Figure S3.** Western blot of TRAIL conjugated vesicles

**Figure S4.** Expanded viability study of soluble TRAIL up to 500 ng/mL

**Figure S5.** Vesicle binding to Jurkat cells

**Figure S6.** Viability of different cell types incubated with unconjugated vesicles

**Figure S7.** Viability of DSPC TRAIL vesicles to different cell lines

**Figure S8.** Caspase 3 and Caspase 8 activity of U-937 cells

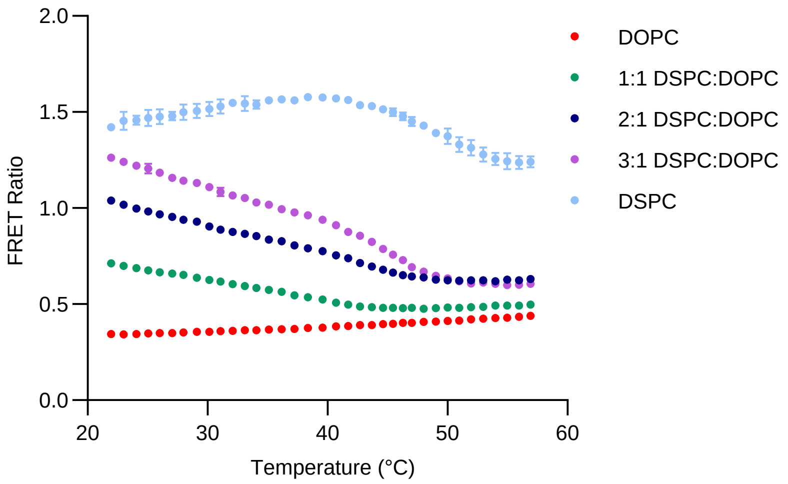


Figure S1. Temperature dependence of lipid domains measured by FRET ratio. Error bars represent SEM from n = 3 different vesicle preparations. Increasing temperatures lead to a convergence of FRET ratios for most lipid compositions, indicating FRET signals reflect membrane domains and temperature induced dissolution of domains leads to similar FRET signals.

Table S2**.** Summary values of size, polydispersity (PDI), and Zeta potential of TRAIL-conjugated vesicles (n = 3 different vesicle preparations).

| | **Size (nm)** | **PDI** | **Zeta Potential (mV)** |
|---|---|---|---|
| **DOPC** | 165 ± 59 | 0.13 | -3.8 ± 1.2 |
| **1:1 DSPC:DOPC** | 161 ± 52 | 0.10 | -4.4 ± 0.8 |
| **2:1 DSPC:DOPC** | 170 ± 57 | 0.11 | -3.7 ± 0.9 |
| **3:1 DSPC:DOPC** | 171 ± 51 | 0.09 | -4.0 ± 0.3 |
| **DSPC** | 191 ± 65 | 0.12 | -3.4 ± 0.9 |

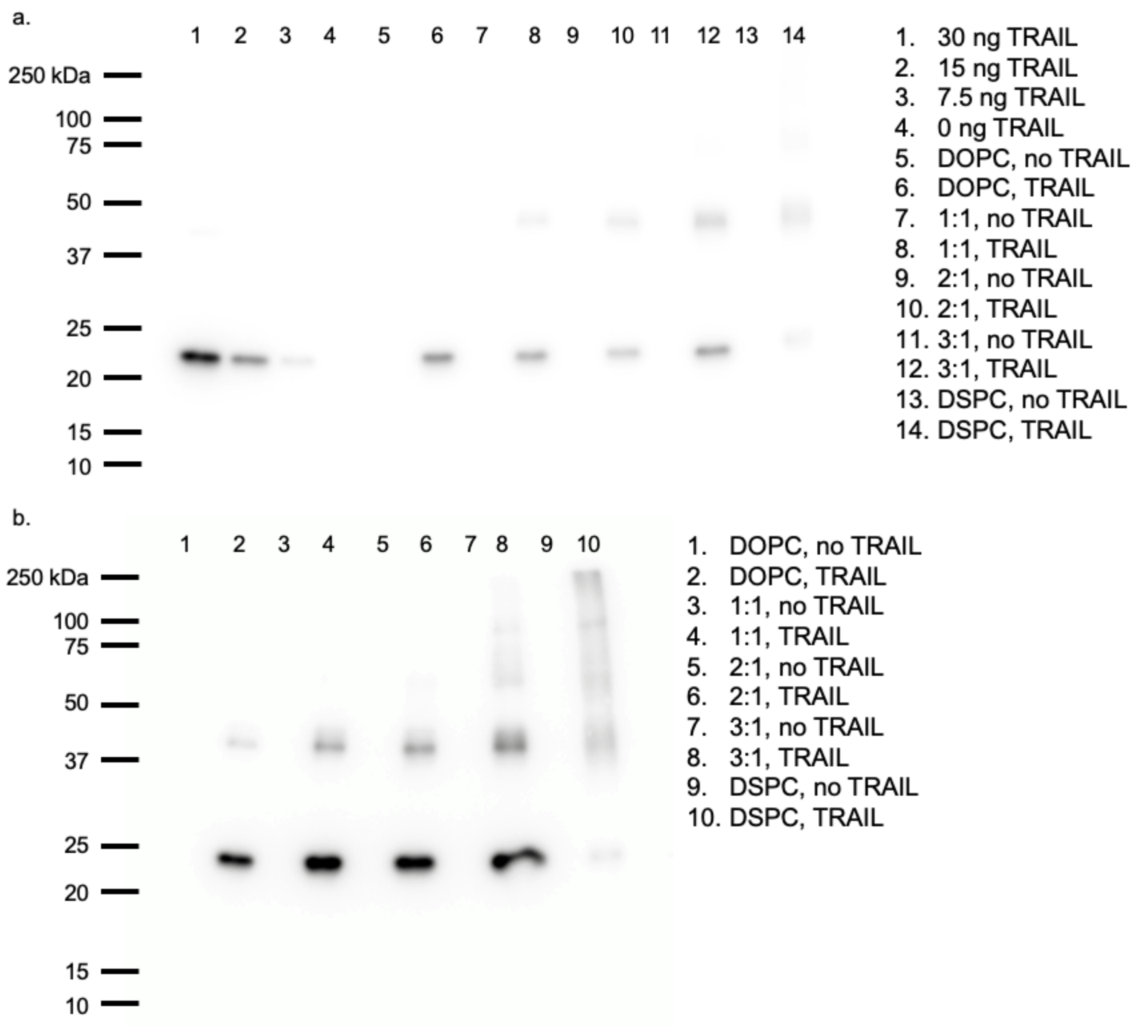


Figure S3. Western blot of TRAIL conjugated vesicles. (a) Representative uncropped anti-TRAIL Western Blot of TRAIL-conjugated vesicles. Corresponding lane key is to the right of the blot and molecular weight markers are labeled in kDa to the left of the blot. A standard curve was generated using purified TRAIL (lanes 1-4) and used to calculate the concentration of TRAIL conjugated to vesicles by densitometry. (b) Representative anti-TRAIL Western Blot performed using non-reducing conditions of TRAIL labeled vesicles. Corresponding lane key is to the right of the blot and molecular weight markers are labeled in kDa to the left of the blot. As TRAIL becomes more concentrated in $L_d$ domains, TRAIL forms more oligomeric structures, as seen on the blot.

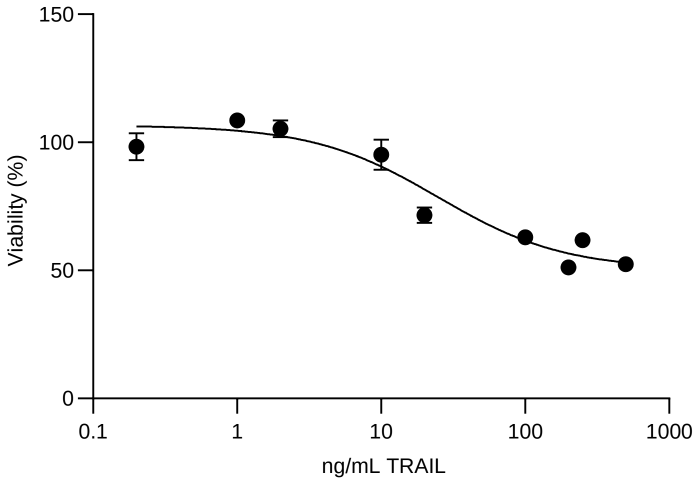


Figure S4: Expanded viability study of soluble TRAIL up to 500 ng/mL. Soluble TRAIL plateaus at approximately 50% viability. Error bars represent SEM from n = 3.

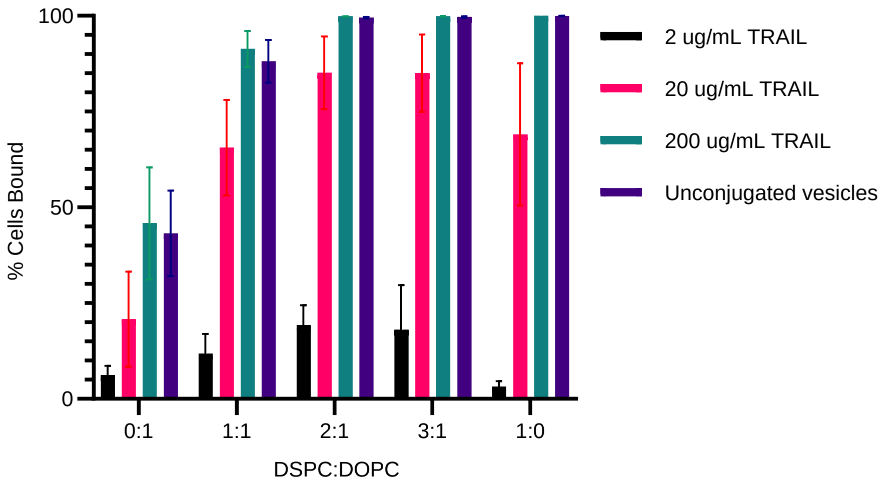


Figure S5. Vesicle binding to Jurkat cells. Unconjugated vesicles are added at the same lipid concentration as 200 µg/mL TRAIL (1 mM lipid concentration) and the amount of cells bound with vesicles is reported. Error bars represent SEM from n = 3 separate flow cytometry experiments with 3 vesicle preparations.

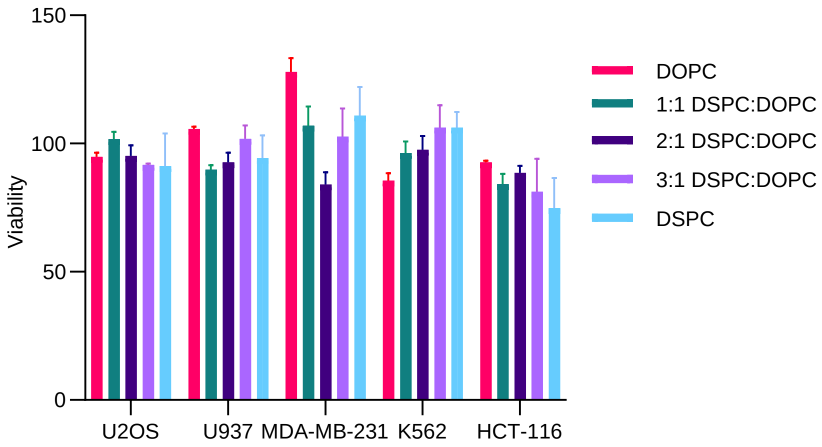


Figure S6. Viability of different cell types incubated with unconjugated vesicles at 1 mM lipid concentration, which corresponds to highest TRAIL concentration tested. Error bars represent SEM from n = 3 different vesicle preparations.

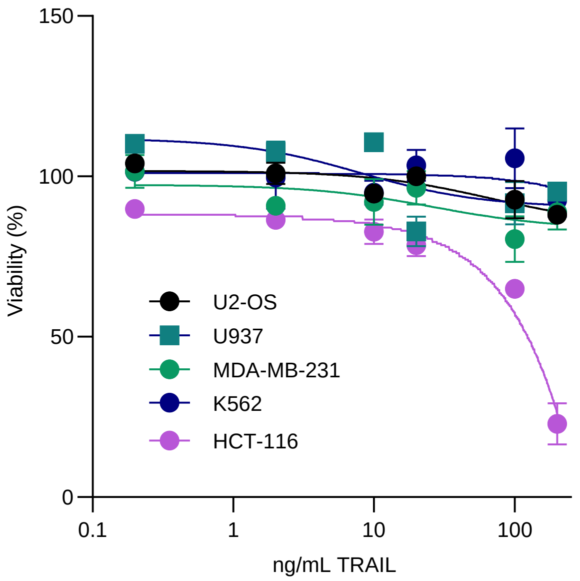


Figure S7. Viability of DSPC TRAIL vesicles to different cell lines. HCT-116 is the only cell line that shows susceptibility to DSPC TRAIL vesicles, while other cell lines do not. Error bars represent SEM from n = 3 different vesicle preparations. Concentration reported is the initial amount of TRAIL added to the vesicles during conjugation.

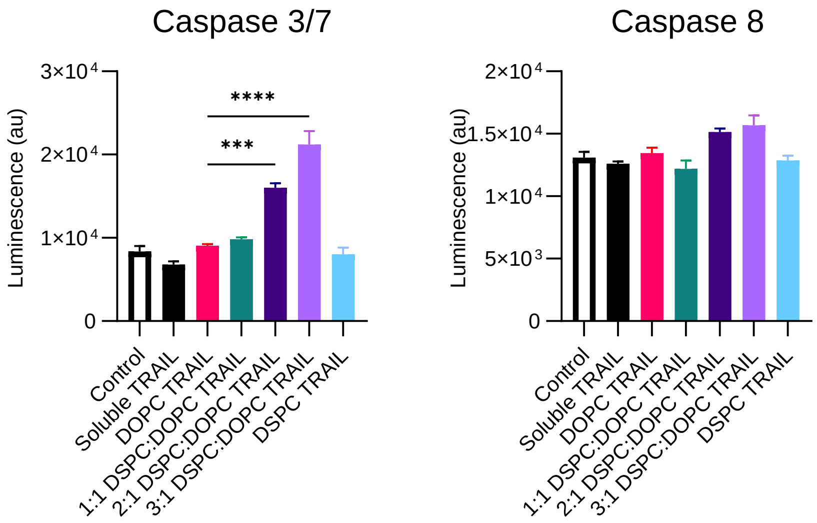


Figure S8: Caspase 3/7 and Caspase 8 activity of U-937 cells incubated with 20 nM TRAIL vesicles and controls. Error bars represent SEM from n = 3 different vesicle preparations. Significance test used was ANOVA. **** $p < 0.0001$, *** $p < 0.001$, ** $p < 0.01$, * $p < 0.05$.
